# Conversion rate to the secondary conformation state in the binding mode of SARS-CoV-2 spike protein to human ACE2 may predict infectivity efficacy of the underlying virus mutant

**DOI:** 10.1101/2021.07.14.452313

**Authors:** Marc M. Sevenich, Joop van den Heuvel, Ian Gering, Jeannine Mohrlüder, Dieter Willbold

## Abstract

Since its outbreak in 2019 SARS-CoV-2 has spread with high transmission efficiency across the world, putting health care as well as economic systems under pressure [1, 2]. During the course of the pandemic, the originally identified SARS-CoV-2 variant has been widely replaced by various mutant versions, which showed enhanced fitness due to increased infection and transmission rates [3, 4]. In order to find an explanation, why SARS-CoV-2 and its emerging mutated versions showed enhanced transfection efficiency as compared to SARS-CoV 2002, an improved binding affinity of the spike protein to human ACE has been proposed by crystal structure analysis and was identified in cell culture models [5-7]. Kinetic analysis of the interaction of various spike protein constructs with the human ACE2 was considered to be best described by a Langmuir based 1:1 stoichiometric interaction. However, we demonstrate in this report that the SARS-CoV-2 spike protein interaction with ACE2 is best described by a two-step interaction, which is defined by an initial binding event followed by a slower secondary rate transition that enhances the stability of the complex by a factor of ∼190 with an overall KD of 0.20 nM. In addition, we show that the secondary rate transition is not only present in SARS-CoV-2 wt but is also found in B.1.1.7 where its transition rate is five-fold increased.

## INTRODUCTION

SARS-CoV-2 is a beta class coronavirus that was first discovered and characterized in Wuhan, China, at the end of 2019 [1, 2]. Since then, it has challenged health care systems due to its rapid spread and COVID-19 transmission throughout the world. Far more than the past coronavirus pandemic, caused by SARS-CoV in 2002, the ongoing pandemic has claimed over 3 Mio lives with over 123 Mio total cases in 221 affected countries, so far [8, 9].

The coronavirus replication depends on a multi-step process starting with the interaction of the viral trimeric spike protein (S) and the human ACE2 receptor that mediates uptake of the viral RNA into the host-cell cytoplasm. Each monomer of the S protein consists of the two functional subunits S1 and S2. The S1 subunit comprises the receptor binding domain (RBD) that interacts via a defined motif sequence (RBM) with an N-terminally located helical structure of the hACE2. The S2 subunit however, plays a crucial role in the membrane fusion process. While the S1-hACE2 interaction allows viral attachment to the host cell surface, the S2’-site is cleaved by the human endoprotease Furin. This leads to irreversible conformational changes of the S protein that result in cell membrane fusion and viral host-cell uptake [10]. Although the viral S proteins of SARS-CoV and SARS-CoV-2 share ∼76 % amino acid sequence identity, SARS-CoV-2 shows an enhanced cell infectivity and human to human transmission efficiency when compared to SARS-CoV [11, 12]. Since its first appearance in 2019, the virus has undergone numerous mutational events, resulting in variants with enhanced fitness concerning their transmissibility [3, 13]. A large proportion of these mutations cluster in the spike protein, where one third of the sequence has been associated with diverse alterations [14]. To date, the most widespread mutant B.1.1.7 has widely replaced the originally identified SARS-CoV-2 virus due to its enhanced fitness [15, 16].

In order to understand why certain variants increase infectivity, the process of viroid contact with the cell surface and cell uptake has been drawn into the center of attention. Although a more efficient fusion process will also impact the infectiousness of the virus [17], the interaction of the spike protein with hACE2 will provide the initial contact and therefore limit the time frame for subsequent processes. Do date, the kinetics of the SARS-CoV spike - hACE2 interaction have been widely defined as a one-step binding process with a mono-exponential decay using a Langmuir based 1:1 fitting model for surface plasmon resonance or biolayer interferometry experiments [5, 7, 10, 18-20]. However, this model fails to describe the complexity of the interaction, which gets apparent looking at the heterogeneity of the complex decay.

Here we report that the interaction of the monomeric SARS-CoV-2 ectodomain with the monomeric hACE2 has a two state mode. The additional secondary conformational transition increases the overall stability of the prefusional state dramatically and therefore enlarges the time frame for initiation of membrane fusion and viral cell entry. In addition, we characterize the secondary interaction state in the trimeric SARS-CoV-2 B.1.1.7 spike protein, where the conversion rate to the secondary conformational state is dramatically increased as compared to the CoV-2 wild type. This observation gives insights in how the infectivity among SARS CoV-2 mutants is modified and represents a precise and fast analysis method to predict the infectivity of novel SARS-CoV-2 variants with mutated spike protein sequences.

## MATERIALS AND METHODS

### Protein expression and purification

#### hACE2

The biotinylated hACE2-fc (residues 18-740) was purchased by Acrobiosystems and contains a C-terminal IgG1 Fc-Tag (residues 100-330), followed by a biotinylated Avitag. Expression was performed in HEK293 cells. The protein has a calculated MW of 111.7 kDa and migrates as 125-150 kDa protein under reducing conditions (SDS-PAGE) due to glycosylation. The monomeric status of the hACE2 protein was verified.

#### Spike protein constructs

The SARS-CoV spike protein constructs were expressed in HEK293-6E cells and purified via a C-terminally introduced 6 X His-Tag with Ni-NTA affinity chromatography. The identity and purity of each construct was verified by SEC-MALS. The foldon sequence of the T4 bacteriophage was C-terminally introduced for the SARS-CoV-2, SARS-CoV 2002 and SARS-CoV-2 B.1.1.7 trimeric constructs in order to promote complex formation [21].

### Kinetic experiments

#### Biolayer interferometry

BLI kinetic experiments were performed with an Octet RED 96 BLI system using streptavidin coated high precision SAX-sensors (ForteBio) and a shaking speed of 1000 rpm. hACE2 was immobilized to a binding level of 1.6 nm with a concentration of 5 µg/ml. Serial dilutions of S1-S2 SARS CoV-2 spike protein were prepared in range of 3.9 to 250 nM in 10 mM HEPES pH 7.4, 150 mM NaCl, 3 mM EDTA, 0.005% Tween-20. The experiment was performed with double reference subtraction.

#### Surface plasmon resonance kinetic experiments

SPR kinetic experiments were performed with a T200-SPR Biacore system (Cytiva) using a Protein A/G coated sensor chip (PAGD-200M, Xantec) and a flow rate of 30 µl/min unless otherwise noted. hACE2 was captured to a response level of 70 RU for each cycle using a concentration of 2.5 µg/ml. The surface was regenerated with 10 mM NaOH by 2x 30 s injections at 10 µl/min. For kinetic measurements serial dilutions of spike protein were prepared in range from 3.9 to 62.5 nM for the monomeric S1S2 and 0.65 to 50 nM for the trimeric constructs in 10 mM HEPES pH 7.4, 150 mM NaCl, 3 mM EDTA, 0.005 % Tween-20. Data fitting was performed using Biacore T200 data evaluation software v. 3.2. For all fits a contribution of refractive index was excluded.

While testing for second state interaction hACE2 was immobilized to a response level of 25 RU for each cycle. 500 nM S1-S2 spike protein was injected with contact times of 150 – 600 s. Buffer referencing was performed prior to each analyte injection cycle. The experimental evaluation was done by alignment of the dissociation start and normalization of the saturation response signal to 100 % by the time point of injection phase end.

## RESULTS AND DISCUSSION

### The Langmuir 1:1 binding model is not sufficient to fit experimental data for spike protein binding to hACE2

For characterization of S1-S2 spike protein interaction with the hACE2 receptor two different kinetic methods were applied. First, for biolayer interferometry hACE2 was immobilized via a C-terminal biotin on a Streptavidin coated sensor-surface. Incubation with a serial dilution of the S1-S2 spike protein yielded the sensograms shown in Fig. 1 A. To obtain apparently good looking fits based on a Langmuir 1:1 model, the dissociation time needed to be significantly shortened (Fig. 1B). Such incomplete fitting allowed the determination of a KD and kinetic rates similar to those that have been published previously [5, 10, 19, 20]. The model fails, however, to describe the dissociation phase of the interaction, which cannot be fitted satisfactorily by a mono-exponential decay (Fig. 1 C).

**Figure 1:**
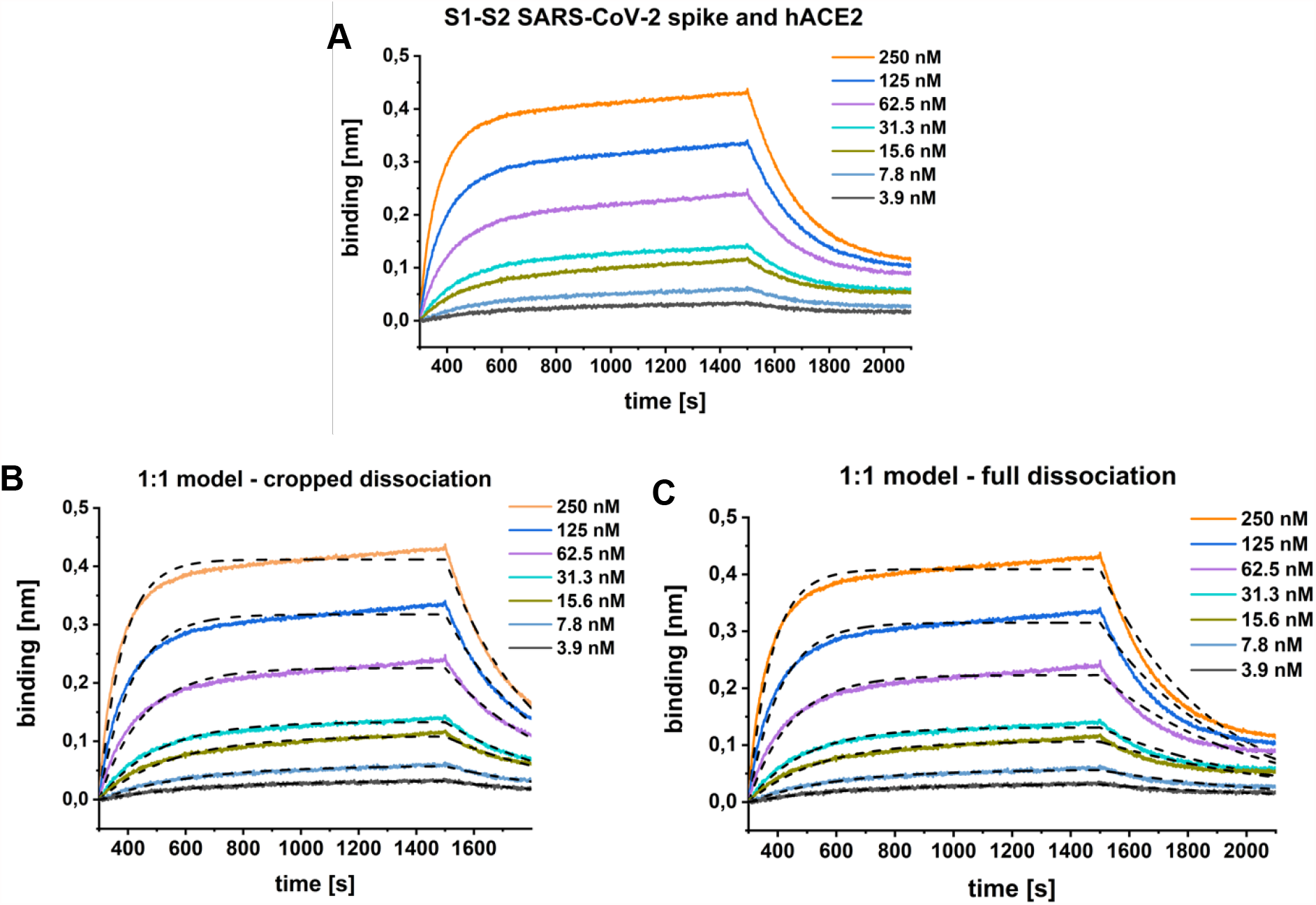
BLI kinetic experiment with SARS-CoV-2 S1-S2 and hACE2. The sensogram (**A**) was globally fitted with a 1:1 interaction model (**B** and **C**, black dashed lines) either with a cropped dissociation time of 200 s (**B**) or the full dissociation time of 600 s (**C**). (**B**) K_D_: 36.3 nM; k_a_: 1.1E+5 +/- 7.5E+4 [1/Ms]; k_d_: 2.4E-3 +/- 4.7E-3 [1/s]. R^2^: 0.98. (**C**) K_D_: 24.2 nM; k_a_: 1.7E+5 +/- 1.5E+5 [1/Ms], k_d_: 1.9E-3 +/- 4.7E-3 [1/s]. R^2^: 0.96.

This conclusion is confirmed by surface plasmon resonance (SPR) experiments. hACE2 was coupled via IgG1 fc-tag on a Protein A/G derivatized surface and various concentrations of S1-S2 spike protein was applied as analyte (Fig 2 A). Apparently satisfying fits based on a Langmuir 1:1 model are only obtained, when the dissociation time is only a few hundred seconds (Fig. 2 B). The inclusion of longer dissociation times into the analysis, however, clearly shows that the interaction of the viral S1-S2 spike protein and the hACE2 is not of a 1:1 Langmuir binding model (Fig. 2 C).

**Figure 2:**
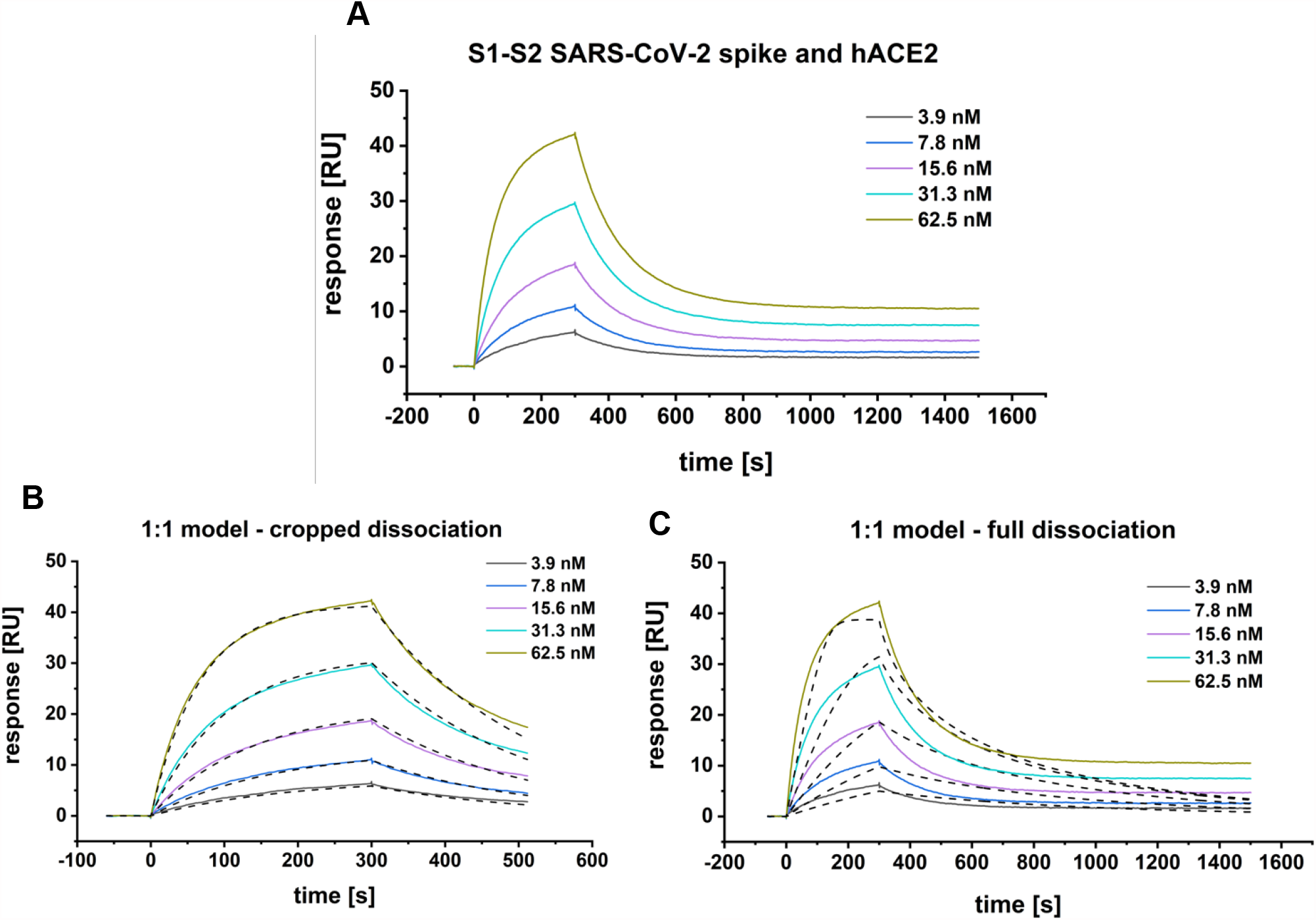
SPR-multicycle kinetic experiment of SARS-CoV-2 S1-S2 and hACE2. The sensogram (**A**) was globally fitted with a Langmuir 1:1 interaction model (**B** and **C**, black dashed lines) either with a cropped dissociation time of 200 s (**B**) or the full (**C**) dissociation time of 1200 s. (**B**) K_D_: 28.5 nM; k_a_ 1.7E+5 +/- 5.4E+2 [1/Ms]; k_d_: 4.7E-3 +/- 1.0E-5 [1/s]. Chi^2^: 0.28 [RU^2^]. (**C**) K_D_: 12.15 nM; k_a_: 5.4E7 +/- 2.2E+6 [1/Ms], k_d_: 0.7 +/- 0.03 [1/s]. Chi^2^: 6.63 [RU^2^].

### S1-S2 spike protein hACE2 interaction induces a time dependent secondary state

Because the interaction of hACE2 and S1-S2 spike protein is not matching a 1:1 Langmuir binding model, we checked for the existence of a potentially underlying second rate process as shown in Fig. 3 [22, 23]. Briefly, hACE2 was coupled at a constant immobilization level of 25 RU. S1-S2 spike protein was injected at a concentration of 500 nM. Contact times were gradually increased by increasing the injection times ranging from 150 to 600 s. For each injection steady-state was reached within a short time interval, so that a constant complex concentration can be assumed during the different contact times.

**Figure 3:**
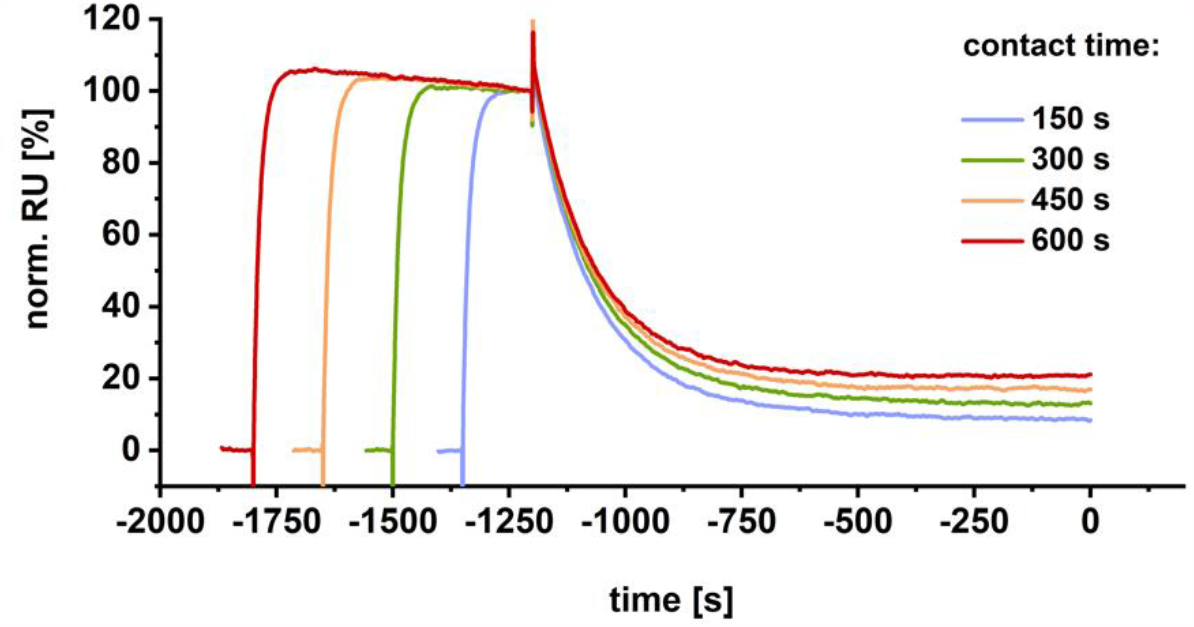
SPR-experiment testing for the presence of a two state reaction. 500 nM SARS-CoV-2 S1-S2 spike protein was injected over a constant immobilization level of 25 RU hACE2 and normalized to 100 % RU for the time point of injection phase end. Injection times were gradually increased (150 – 600 s).

The resulting plot reveals a strong dependency of incubation time and dissociation rate (Fig. 3). Increasing injection times and thus increasing contact time correlates with decreasing dissociation rates. This finding cannot be expected for a single step 1:1 interaction, but clearly indicates the formation of a secondary complex state, whose proportion increases with contact time duration. This is typical for a two-step binding mode, in which the formation of the primary complex induces a conformational reorganization into a secondary complex conformation that strengthens the interaction and leads to a very slow dissociation rate.

### Secondary complex state of S1-S2 spike protein with hACE2 results in enhanced complex stability

In order to define the secondary rate kinetics of complex transition after primary binding, the interaction of S1-S2 spike protein and hACE2 was fitted with a secondary state model. In contrast to the previously presented attempts of the Langmuir model fitting (Fig. 1 and 2), the two step kinetic model allows the description of the interaction with high accuracy over the complete dissociation time (Fig. 4 A).

**Figure 4:**
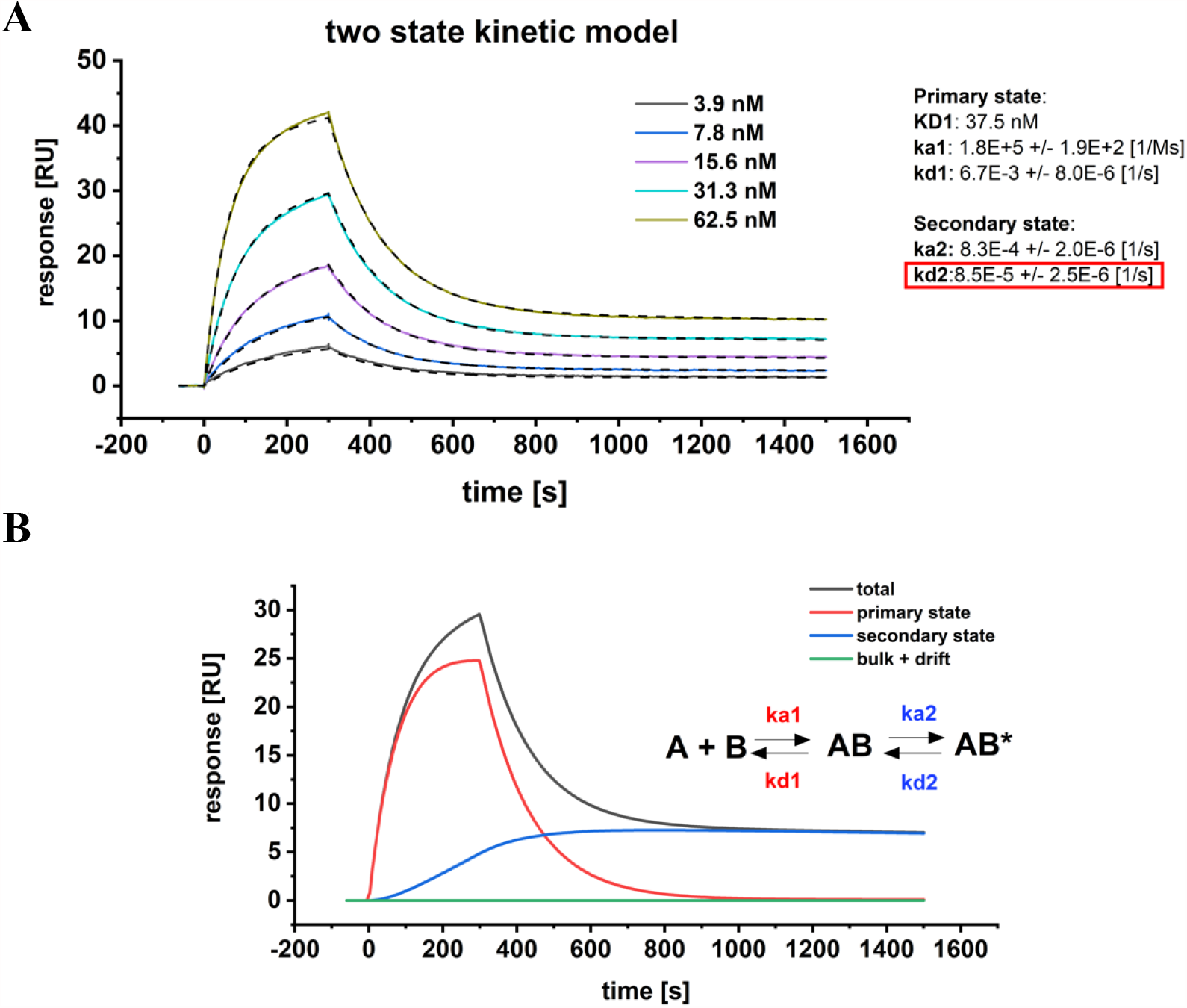
SPR-multicycle kinetic experiment of SARS-CoV-2 S1-S2 and hACE2. **(A)** The sensogram was globally fitted with a two state kinetic model including the full dissociation time of 1200 s. Kinetic parameters for the first interaction were determined with K_D_1 of 37.5 nM, k_a1_ of 1.8E+5 +/- 1.9E+2 [1/Ms] and k_d1_ of 6.6E-3 +/- 8.0E-6 [1/s]. Kinetic parameters for the second interaction were determined with k_a2_ of 8.3E-4 +/- 2.0E-6 [1/s] and k_d2_ of 8.54E-5 +/- 2.5E-6 [1/s]. Chi^2^: 0.08 [RU^2^]. The overall K_D-total_ for both events was identified with 0.2 nM. **(B)** Component analysis of the 31.3 nM S1-S2 spike protein binding curve as shown in (A). The sensogram (total) is composed of the primary binding event (red), followed by a secondary transition event (blue) which results in a highly stable secondary complex (AB*).

The kinetic rates of the primary binding event were identified with ka1 of 1.8 · 10^−5^ M^-1^ s^-1^ and kd1 of 6.7 · 10^−3^ s^-1^ resulting in a KD of 37.5 nM. This matches with previously reported kinetic values for the RBD interaction with ACE2 [5, 10, 19, 20]. As soon as the first complex is formed, a secondary event increases the complex stability (Fig. 4 B). This transition is most likely a structural rearrangement that decreases the complex dynamics as proposed previously [6]. The kinetic data show that this secondary process is with an on-rate of 8.3 · 10^−4^ s^-1^ rather slow compared to the primary binding event, but at the same time increases the complex half-life by a factor of ∼80 with an off-rate of 8.5 · 10^−5^ s^-1^. When the kinetic values of the primary and secondary events are combined to one binding constant, the full binding process yields a total affinity that is described with KD-total of 0.20 nM.

### The SARS-CoV-2 RBD interaction with hACE2 follows a Langmuir interaction kinetic

To verify whether the secondary transition can be exclusively obtained for the S1S2 monomeric construct or might be associated with certain proportions of the spike protein, a SARS-CoV-2 RBD construct was assayed for the existence of an underlying secondary rate kinetic.

When the SARS-CoV-2 RBD was analyzed in a multicycle kinetic experiment on the immobilized hACE2, the dissociation phase showed a clear mono-exponential behavior with complete baseline dissociation, which is in full agreement with a Langmuir based 1:1 interaction model (Fig. 5 A). This finding appears to be in contrast with the previously identified biphasic dissociation for the S1S2 CoV-2 construct. Additionally, the RBD did not show a contact time dependent alteration of the dissociation phase, when increasing contact times are applied (Figure 5 B). However, the dissociation constant as well as on- and off-rate of the RBD and the primary binding event of the monomeric S1S2 show high similarity. This implies that the primary binding event of the two-state interaction is carried out by the interaction of the RBD and hACE2 alone, whereas the context of the full S1S2 protein is essential for the formation of the secondary complex.

**Figure 5:**
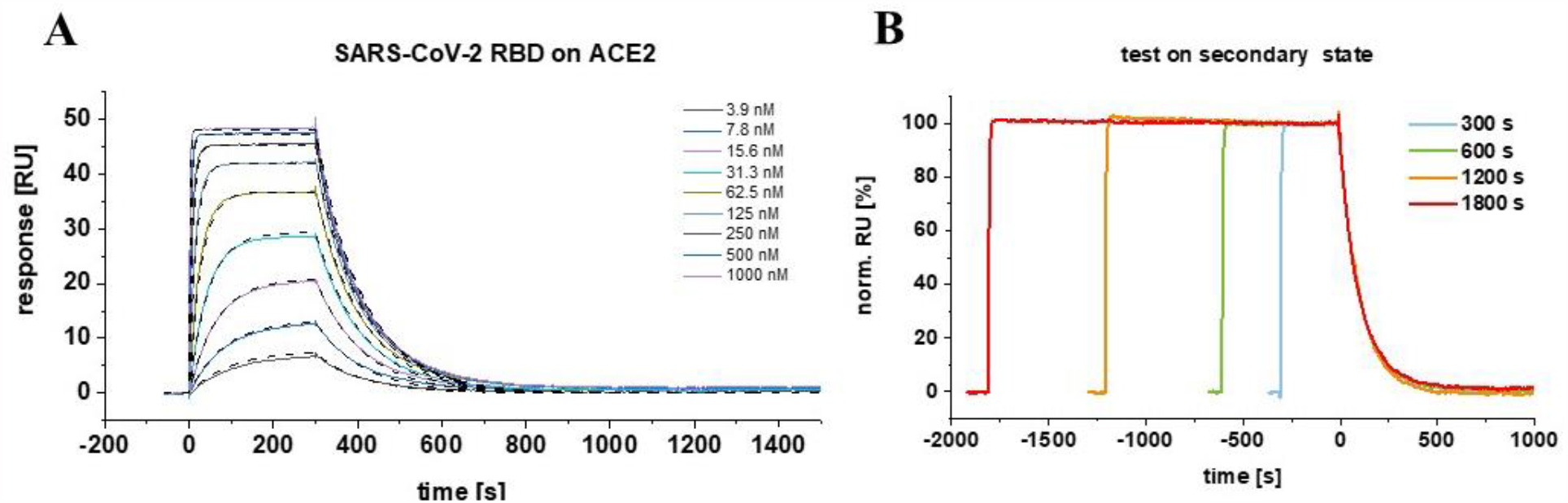
The SARS-CoV-2 RBD interaction with hACE2 follows a Langmuir based kinetic with time-independent mono-exponential decay. **(A)** Multicycle experiment with SARS-CoV-2 RBD. hACE2-fc was immobilized on a protein A/G sensor chip and SARS-CoV-2 RBD was injected in concentration range of 3.9 – 1000 nM. The K_D_ was globally fitted with a 1:1 Langmuir based interaction model. The kinetic parameters were determined with K_D_ of 21.3 nM, k_a_ of 4.25 E+5 +/- 2.2 E+2 [1/Ms] and k_d_ of 9.1 E-3 +/- 4.2E-6 [1/s]. (**B**) Test on secondary state reaction. hACE2-fc was immobilized on a protein A sensor surface and 500 nM of SARS-CoV-2 was injected at a constant concentration for increasing contact intervals. Time points of injection phase end were normalized to 100 % and the dissociation starting point was aligned on the time scale.

### The secondary state transition rate is modified among SARS-CoV, SARS-CoV-2 and B.1.1.7 trimeric spike proteins

The secondary state model was essential to fully describe the interaction of the S1S2 monomer with hACE2. However, the physiological quarternary structure of the spike protein is a homotrimer, where the interaction with hACE2 is mediated by a single RBD in its “up-conformation”. During this first contact the other two RBD’s remain in their closed state, which does not allow direct contact with hACE2 [24]. Hence, the interaction of the trimeric spike protein and hACE2 is defined by a 1:1 stoichiometry.

Next, we analyzed the interaction of the SARS-CoV 2002, SARS-CoV-2 wt and SARS-CoV-2 B.1.1.7 mutant trimeric spike protein with hACE2 as described previously. The sensograms were again fitted using global fitting with the two state reaction model.

The dissociation phase of the trimeric spike proteins (Fig. 6A) shows a biphasic decay as already observed for the monomeric SARS-CoV-2 S1S2. Again, the secondary state model allowed the best fit for the given sensograms. The component analysis reveals that the secondary state transition (Fig. 6A, blue line) is dominating the complex formation after short initial contact time. Figure 6 B shows the on-off chart of the primary and secondary interaction mode as fitted for the SARS-CoV 2002, the SARS-CoV-2 wt and SARS-CoV-2 B1.1.7 trimers. When the here identified ka and kd values of the primary interaction are compared with those found for the SARS-CoV-2 S1S2 and RBD constructs, the 2002 and wt trimeric spikes show values in the same range, implying that the initial binding event is not so much different among these constructs. The B.1.1.7 mutant however, shows a faster association and dissociation for the primary contact. Similarly, the kinetic values of the B.1.1.7 secondary state deviate from those identified for the 2002 and wt trimers. Here, the secondary state transition rate is of highest interest, because it will directly impact the contact time that will be needed to form a complex with enhanced stability. The ka identified for B.1.1.7 is 5.1 and 6.1 faster than the transition rate for the wt and 2002 trimer, respectively.

**Figure 6:**
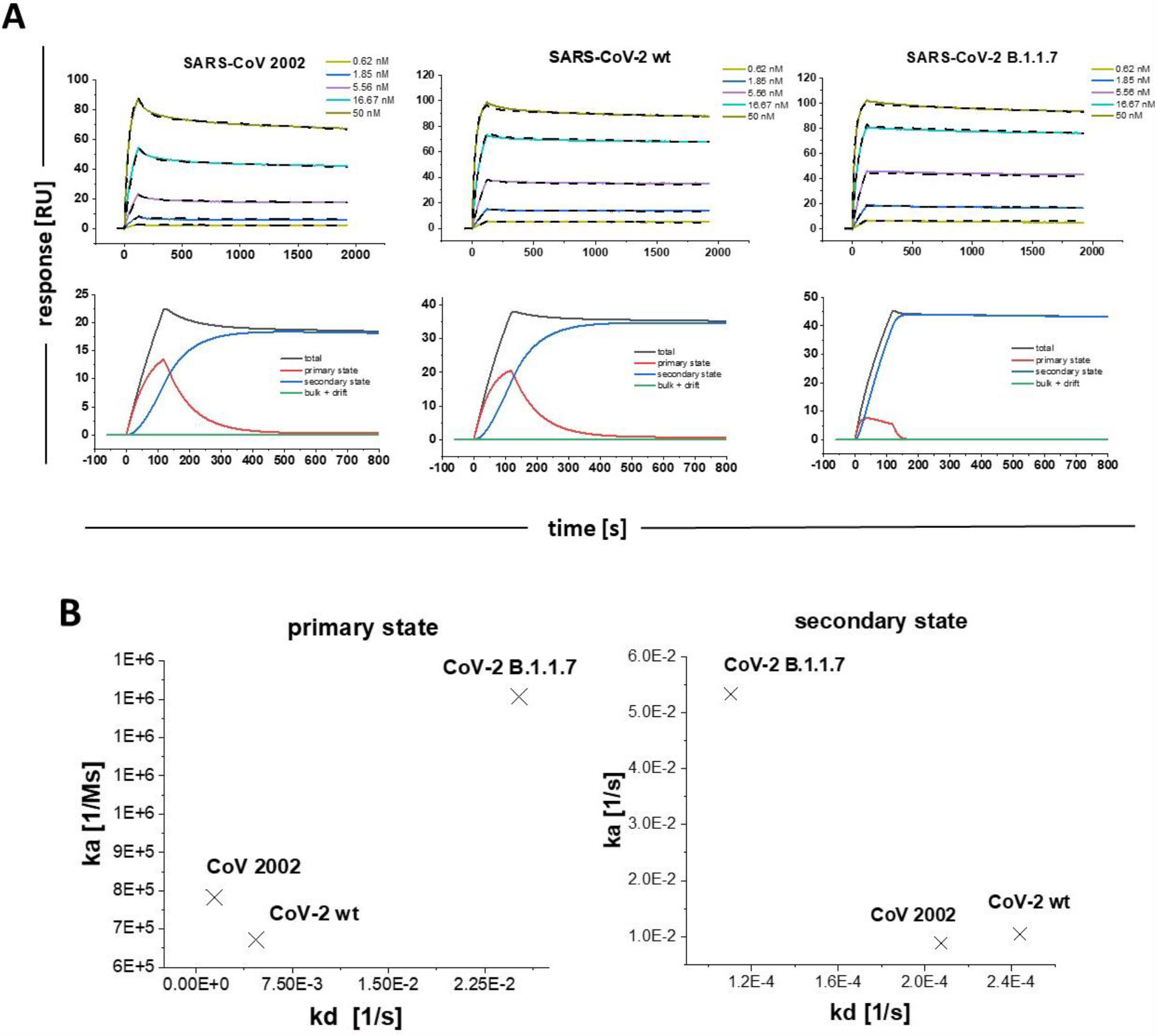
Multi cycle SPR experiments with SARS-CoV 2002, SARS-CoV-2 wt and SARS-CoV-2 B.1.1.7 trimeric spike proteins. **(A)** hACE2-fc was immobilized on a protein A/G sensor surface and CoV trimer proteins were injected in concentration range of 0.62 – 50 nM. The sensograms were globally fitted with the secondary state reaction model. **(SARS-CoV 2002)**: K_D1_ 6.7 nM, k_a1_ 6.72 E+5 ± 4.4E+3 [1/Ms], k_d1_ 4.68 E-3 [1/s] ± 6.9E-5, k_a2_ 8.80 E-3 ± 6.9E-5 [1/Ms], k_d2_ 2.07E-4 ± 1.60 E-6 [1/s]; K_D-total_ 160 pM **(SARS-CoV-2 wt)**: K_D1_ 1.8 nM, k_a1_ 7.82 E+5 ± 1.2E+3 [1/Ms], k_d1_ 1.41 E-3 ± 3.75E-5 [1/s], k_a2_ 1.05E-2 ± 1.8E-4 [1/Ms], k_d2_ 2.44E-4 ± 3.5E-6 [1/s]; K_D-total_ 41 pM. (**SARS-CoV-2 B.1.1.7**) K_D1_ 192.7 nM, k_a1_ 1.31 E+6 ± 1.5E+4 [1/Ms], k_d1_ 2.5 E-2 ± 1.5E-3 [1/s], k_a2_ 5.3E-2 ± 1.5E-3 [1/Ms], k_d2_ 1.1 E-4 ± 2.6E-6 [1/s]; K_D-total_ 40 pM. For graphic representation of the distribution of primary and secondary state reaction, a component analysis was performed for each of the trimeric spike proteins using the injection concentration of 5.56 nM spike. The sensogram (total) is composed of the primary binding event (red), followed by a secondary transition event (blue) which results in a highly stable secondary complex (**B**) On–off chart for the kinetic values of the SARS-CoV spike trimers in the primary and secondary state.

The significance of this finding becomes obvious when it is transferred to physiological conditions, where the probability of a potential primary contact between the viroid spike and the cell surface located hACE2 is limited by the local concentration of the two interaction partners. Hence, a more rapid transition of a low to high affinity binding state will increase the complex half-time once a primary contact occurs and therefor increase the infection efficacy as observed for the B.1.1.7 and other mutants.

## SUMMARY AND CONCLUSION

This study helps to understand the basis of the enhanced infectivity that has been observed for SARS-CoV-2 and its derived mutants as compared to SARS-CoV 2002. We have demonstrated that a secondary state of the SARS-CoV-2 spike – hACE2 complex exists and that the transition increases the stability of the complex by a factor of ∼190 with an overall KD of 0.20 nM for the monomeric spike S1S2 protein. Furthermore, when the isolated RBD is tested for its affinity to hACE2, no secondary state formation was observed. This suggests that the context of the whole ectodomain is needed to promote the secondary state within the complex after the first contact is mediated by the RBD. When the kinetics of the monomeric SARS-CoV-2 S1S2 are compared with the trimeric variant, an increased secondary state transition was identified for the trimeric protein, suggesting cooperative effects between the subunits. Finally, the secondary state formation was verified for the trimeric constructs of SARS-CoV 2002 and SARS-CoV-2 B.1.1.7, where a 5 to 6 fold increased secondary state transition rate was observed for the B.1.1.7 mutant compared to the wt and 2002 trimeric spike protein.

Taken together, these findings highlight the role of the SARS-CoV-2 spike protein in the context of the ongoing pandemic and stress its importance as potential drug-target. The presented SPR method for the verification of secondary state transitions within the spike protein – hACE2 complex allows a straightforward way of predicting the infectiousness of novel SARS-CoV-2 mutants. To date, several studies aim for the inhibition of the spike – hACE2 interaction by targeting one of the interaction partners [25-28]. However, the present study shows that an efficient inhibitor should impact both, the primary as well as secondary binding state in order to obtain significant reduction of complex formation. Since the secondary transition rate is potentially the defining parameter for an enhanced infection rate, it is important to understand its molecular mechanism. Targeting the secondary state transition could represent a very efficient therapeutic mechanism for the therapy of COVID-19. Most importantly, we suggest to use the full quantitative kinetic and thermodynamic binding behavior of newly appearing spike protein variants to hACE2 to possibly predict the infectivity efficacy of the underlying virus mutation.

## AUTHOR CONTRIBUTIONS

D.W. and J.M. generated the project strategy. M.M.S. planned and performed the SPR and BLI experiments and conducted the data evaluation. J.V.D.H. provided the purified and characterized S1-S2 spike protein. I.G. contributed to the experimental design. M.M.S and D.W. took primary responsibility for writing the manuscript. All authors edited the manuscript

## REFERENCES

1. Huang, C., et al., Clinical features of patients infected with 2019 novel coronavirus in Wuhan, China. Lancet, 2020. 395(10223): p. 497–506.

2. Li, Q., et al., Early Transmission Dynamics in Wuhan, China, of Novel Coronavirus-Infected Pneumonia. N Engl J Med, 2020. 382(13): p. 1199–1207.

3. Chen, J.H., et al., Mutations Strengthened SARS-CoV-2 Infectivity. Journal of Molecular Biology, 2020. 432(19): p. 5212–5226.

4. Plante, J.A., et al., Spike mutation D614G alters SARS-CoV-2 fitness. Nature, 2020.

5. Shang, J., et al., Structural basis of receptor recognition by SARS-CoV-2. Nature, 2020. 581(7807): p. 221–224.

6. Brielle, E.S., D. Schneidman-Duhovny, and M. Linial, The SARS-CoV-2 Exerts a Distinctive Strategy for Interacting with the ACE2 Human Receptor. Viruses, 2020. 12(5).

7. Ozono, S., et al., SARS-CoV-2 D614G spike mutation increases entry efficiency with enhanced ACE2-binding affinity. Nature Communications, 2021. 12(1).

8. Rabaan, A.A., et al., SARS-CoV-2, SARS-CoV, and MERS-COV: A comparative overview. Infez Med, 2020. 28(2): p. 174–184.

9. Lee, N., et al., A major outbreak of severe acute respiratory syndrome in Hong Kong. N Engl J Med, 2003. 348(20): p. 1986–94.

10. Wang, Q., et al., Structural and Functional Basis of SARS-CoV-2 Entry by Using Human ACE2. Cell, 2020. 181(4): p. 894–904 e9.

11. Hoffmann, M., et al., SARS-CoV-2 Cell Entry Depends on ACE2 and TMPRSS2 and Is Blocked by a Clinically Proven Protease Inhibitor. Cell, 2020. 181(2): p. 271–280 e8.

12. Xia, S., et al., Inhibition of SARS-CoV-2 (previously 2019-nCoV) infection by a highly potent pan-coronavirus fusion inhibitor targeting its spike protein that harbors a high capacity to mediate membrane fusion. Cell Res, 2020. 30(4): p. 343–355.

13. Tang, J.T.P.H. D, Emergence of a new SARS-CoV-2 variant in the UK. Journal of Infection, 2021. 82(4): p. e27–e28.

14. Guruprasad, L., Human SARS CoV-2 spike protein mutations. Proteins-Structure Function and Bioinformatics, 2021. 89(5): p. 569–576.

15. Tang, J.L.W., P.A. Tambyah, and D.S.C. Hui, Emergence of a new SARS-CoV-2 variant in the UK. Journal of Infection, 2021. 82(4): p. E27–E28.

16. Piantham, C. and K. Ito, Estimating the increased transmissibility of the B.1.1.7 strain over previously circulating strains in England using fractions of GISAID sequences and the distribution of serial intervals. medRxiv, 2021.

17. Coutard, B., et al., The spike glycoprotein of the new coronavirus 2019-nCoV contains a furin-like cleavage site absent in CoV of the same clade. Antiviral Res, 2020. 176: p. 104742.

18. Wang, Y., M. Liu, and J. Gao, Enhanced receptor binding of SARS-CoV-2 through networks of hydrogen-bonding and hydrophobic interactions. Proc Natl Acad Sci U S A, 2020. 117(25): p. 13967–13974.

19. Wrapp, D., et al., Cryo-EM structure of the 2019-nCoV spike in the prefusion conformation. Science, 2020. 367(6483): p. 1260–1263.

20. Lan, J., et al., Structure of the SARS-CoV-2 spike receptor-binding domain bound to the ACE2 receptor. Nature, 2020. 581(7807): p. 215–220.

21. Papanikolopoulou, K., et al., Formation of highly stable chimeric trimers by fusion of an adenovirus fiber shaft fragment with the foldon domain of bacteriophage T4 fibritin. Journal of Biological Chemistry, 2004. 279(10): p. 8991–8998.

22. H Saunal, R.K., MHV Van Regenmortel Antibody affinity measurement. Immunochemistry, 1997

23. Lund-Katz, S., et al., Surface plasmon resonance analysis of the mechanism of binding of apoA-I to high density lipoprotein particles. J Lipid Res, 2010. 51(3): p. 606–17.

24. Gur, M., et al., Conformational transition of SARS-CoV-2 spike glycoprotein between its closed and open states. Journal of Chemical Physics, 2020. 153(7).

25. Alagumuthu, M., S. Rajpoot, and M.S. Baig, Structure-Based Design of Novel Peptidomimetics Targeting the SARS-CoV-2 Spike Protein. Cellular and Molecular Bioengineering, 2021. 14(2): p. 177–185.

26. Han, Y.X. and P. Kral, Computational Design of ACE2-Based Peptide Inhibitors of SARS-CoV-2. Acs Nano, 2020. 14(4): p. 5143–5147.

27. Guo, L., et al., Engineered trimeric ACE2 binds viral spike protein and locks it in “Three-up” conformation to potently inhibit SARS-CoV-2 infection. Cell Research, 2021. 31(1): p. 98–100.

28. Bojadzic, D., O. Alcazar, and P. Buchwald, Methylene Blue Inhibits the SARS-CoV-2 Spike-ACE2 Protein-Protein Interaction-a Mechanism that can Contribute to its Antiviral Activity Against COVID-19. Frontiers in Pharmacology, 2021. 11.

